# FERONIA defines intact tissue boundaries through cuticle development

**DOI:** 10.1101/2025.06.09.658546

**Authors:** Gayeon Kim, Jeongho Choi, Ryeo Jin Kim, Eunkyoo Oh, Seung Yong Shin, Hyun-Soon Kim, Hye Sun Cho, Mi Chung Suh, Hyo-Jun Lee

## Abstract

Plant cuticle is the first hydrophobic barrier between the epidermis and the environment. Upon wounding, damaged tissues undergo healing processes that involve cuticle or callus formation at the wound site. However, signaling pathways that initiate cuticle development and callus formation in the wound-proximal region are still poorly understood. Here, we reveal that the FERONIA receptor-like kinase facilitates cuticle development in the epidermis and FER-mediated cuticle formation limits the propagation of wound-induced reactive oxygen species (ROS), which trigger callus formation. Cuticle defects stimulate NADPH oxidase-dependent ROS production, which leads to unrestricted callus formation. However, the cuticle formed in mesophyll cells in the vicinity of the wound suppresses ROS propagation, thereby preventing unorganized callus formation beyond the wound-proximal site and activating programmed cell death adjacent to the wound. These findings provide valuable insights into cuticle development in aerial tissues and its defensive function for preserving the integrity of undamaged regions.

## Introduction

The plant cuticle is a hydrophobic barrier which covers the outermost surface of aerial organs including leaves, stems and flowers^1^. It is primarily composed of polyester cutin and organic solvent-soluble cuticular waxes, which together ensure appropriate organ development and confer resistance against biotic and abiotic stresses^2,3^. The hydrophobicity underlying these critical functions arises from its chemical composition: waxes consist of very-long-chain (C>18, mainly C24-33) fatty acids (VLCFAs), aldehydes, alkanes, ketones, primary and secondary alcohols, and wax-esters, while cutin is made up of C16 and C18 unsubstituted and various substituted fatty acids including dicarboxylic acids and ω-hydroxyacids^4^. Extensive studies have established cuticle’s function as a developmental surface barrier in aerial organs, root caps, and embryos^5–7^.

Beyond its role as a protective surface boundary in developmental processes, cuticle also contributes to tissue repair. Upon wounding or organ abscission, *de novo* cuticle is deposited at the exposed surface, which accompanies transdifferentiation of non-epidermal cells to epidermal cells. Secession cells in the abscission zone (AZ) of the separating organs form lignin ‘braces’, while residuum cells in the AZ of the remaining organs undergo cuticle sealing at the newly formed surface^8^. When wounding induces jasmonic acid-mediated cuticle formation in leaves, wound- and methyl jasmonate-inducible cytochrome P450 plays a role in maintaining cuticle composition and permeability^9,10^. These findings suggest that the cuticle development was induced to fulfill its protective role as the exposed cells became vulnerable to external stressors^10^. While little is known about the molecular mechanisms underlying induction of cuticle initiation or reinforcement, it was only recent when a bidirectional signaling pathway involving ABNORMAL LEAF SHAPE1 (ALE1) subtilase and GASSHO (GSO) receptor-like kinases (RLKs) between embryo and endosperm was reported to facilitate the formation of the embryonic cuticle, which ensures embryo development and survival^7^. Upstream regulation of cuticle barrier after wounding or organ abscission, or during organ expansion after germination, when the mesophyll or epidermal cells are exposed to the terrestrial atmosphere, still has much to be elucidated^11^.

Damaged tissues reprogram the identity of cells specifically near the wound region to preserve organismal integrity or to generate new organs^12^. Callus is a distinct tissue that forms as part of the wound-healing process, primarily through the actions of the phytohormones auxin and cytokinins^13^. Since callus cells can acquire pluripotency, they are capable of regenerating new organs such as roots and shoots^12^. Consequently, callus formation and subsequent organogenesis have been widely utilized in biotechnology-not only to propagate genetically identical individuals but also to generate genetically engineered plants from transgenic callus cells^14^. The molecular mechanisms underlying callus formation have been shown to resemble those of root development. The auxin-responsive transcription factor WUSCHEL-RELATED HOMEOBOX 11 (WOX11) and its downstream target LATERAL ORGAN BOUNDARIES DOMAIN 16 (LBD16) play central roles in this process. Ectopic overexpression of these transcription factors not only promotes callus proliferation but also initiates callus formation throughout the plant body^15,16^, indicating that upregulation of these transcription factors is sufficient to initiate callus formation.

While callus formation occurs throughout the entire root region, it is primarily restricted to wound-proximal areas in the shoots of various plant species, including both monocots and dicots^16–18^. These observations suggest that wound signals are involved in initiating callus formation in shoots. Previous studies have sought to identify the molecular nature of wound signals related to callus formation. Physical damage induces the production of the peptide elicitor REGENERATION FACTOR 1 (REF1), which activates a downstream receptor kinase to upregulate the AP2/ERF transcription factor WOUND-INDUCED DEDIFFERENTIATION 1^19^. However, although treatment with synthetic REF1 peptide promoted callus formation in wound-proximal regions, it was less effective in inducing callus formation at wound-distal regions of tomato hypocotyls^19^, suggesting that REF1 is not the sole wound-derived signal required for callus initiation. Recent studies have provided insights into spatial regulation of callus formation. Glutamate receptor-like proteins act as wound-responsive ion channels, and their inhibition by antagonists expands the callus-forming region beyond the wound-proximal area^20^. Submergence-induced ethylene signaling activates callus formation in wound-distal cells^21^. Nevertheless, the precise molecular mechanisms by which plants distinguish between wound-proximal and -distal regions to regulate callus formation remain poorly understood. Therefore, in this study, we investigated the mechanisms by which a FERONIA (FER) RLK mediates leaf cuticle formation and how newly formed or reinforced cuticle in mesophyll cells near the wound site specifies the site of callus formation and maintains the integrity of undamaged regions.

## Results

### Cuticle defects produce ROS for callus formation

Since callus formation occurred exclusively near the wound site in Arabidopsis leaf explants, we hypothesized that wounding generates signals that initiate callus formation. Among the previously identified wound-induced signals, we focused on reactive oxygen species (ROS), due to their pivotal role in root regeneration from leaf explants, as demonstrated in our previous study^22^. To test our hypothesis, we applied the NADPH oxidase inhibitor diphenyleneiodonium (DPI) to suppress ROS production during callus formation. DPI treatment increased the proportion of leaf explants that failed to form callus and reduced the average callus weight (**Fig. 1a,b** and **Extended Data Fig. 1a**). These data suggest that NADPH oxidase-mediated ROS production is required for callus formation. However, double mutants of *RBOHD* and *RBOHF*, which encode NADPH oxidases, did not show altered callus formation (**Extended Data Fig. 1b**), implying that multiple NADPH oxidases may function redundantly to produce ROS after wounding. When GUS expression and activity were measured in transgenic plants expressing GUS driven by the *WOX11* or *CYCLIN B1;1* (*CYCB1;1*) promoters, we noticed that the expression of marker genes for callus formation (*WOX11*) and cell division (*CYCB1;1*) was suppressed by DPI, further supporting the role of ROS in callus induction (**Fig. 1c,d** and **Extended Data Fig. 1c**). Staining with nitroblue tetrazolium (NBT) and 3,3’-diaminobenzidine (DAB) revealed that both superoxide and hydrogen peroxide accumulated specifically near the wound site, particularly along the vascular tissues (VT) and at the border of callus formation (BC), but DPI treatment notably reduced ROS accumulation at the VT (**Fig. 1e** and **Extended Data Fig. 1d**). These results indicate that wounding induces ROS production via NADPH oxidases and that this ROS production is required for callus formation in leaf explants.

**Fig. 1.**
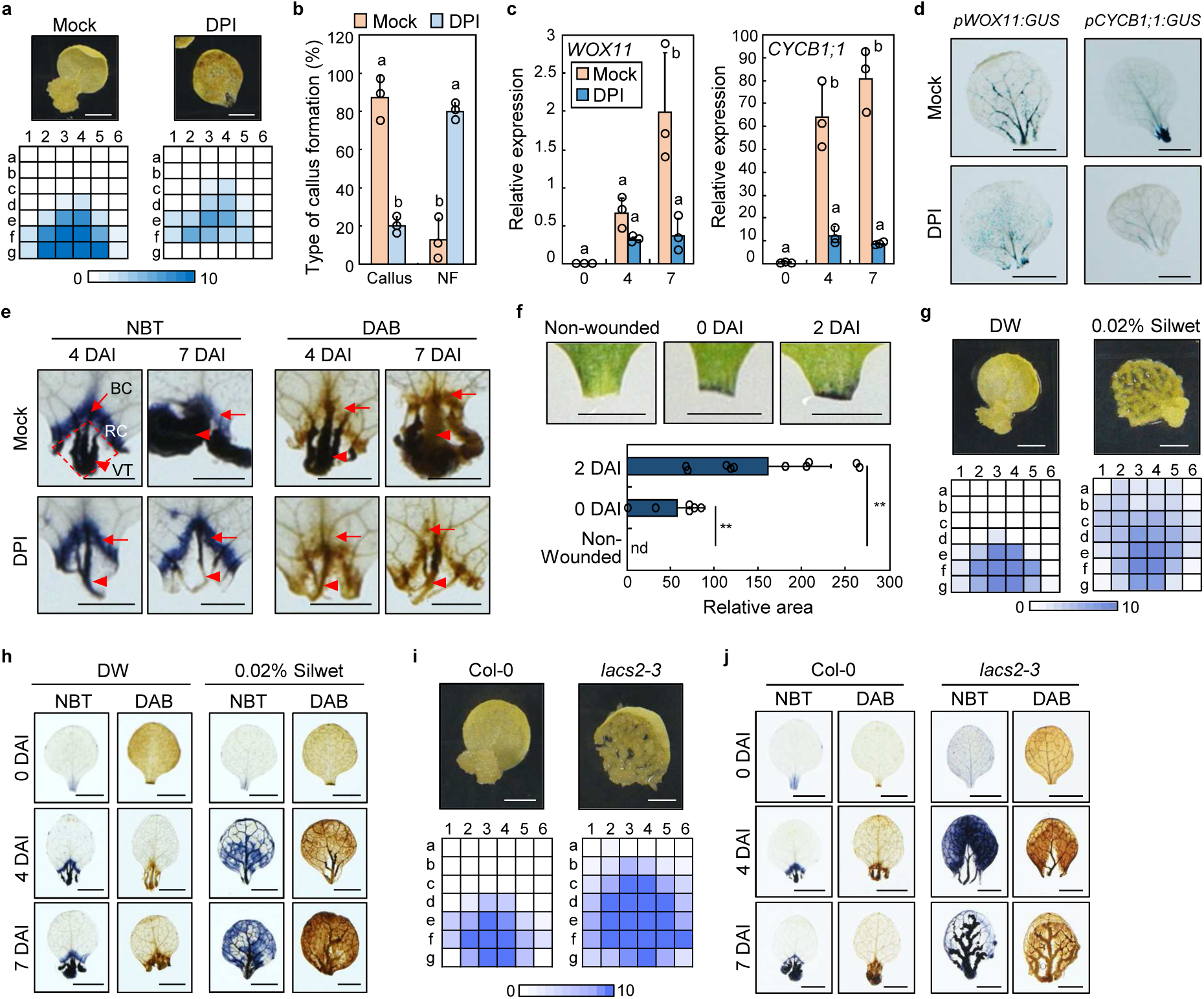
Cuticle defects at wound sites trigger ROS production to induce callus formation. **a**,**b**, Callus formation in Col-0 with or without DPI. Heatmap of callus-forming sites (**a**) and percentage of callus-forming (callus) and non-forming (NF) explants (**b**). Data represent mean ± s.d. (Tukey’s test, *P* < 0.05; *n* = 3). **c**, Effects of DPI on *WOX11* and *CYCB1;1* expression in Col-0. Data represent mean ± s.d. from three independent biological replicates (Tukey’s test, *P* < 0.05). X-axis indicates days after incubation (DAI). **d**, Promoter activity of *WOX11* and *CYCB1;1* by GUS staining at 4 DAI. **e**, NBT and DAB staining in Col-0. Arrows, border of callus formation (BC); arrowheads, vascular tissue (VT); dotted area, region of callus formation (RC). **f**, TBO staining in Col-0. Intact seedlings were stained for non-wounded controls. Other explants were incubated on CIM before staining. TBO signal area was quantified. Data represent mean ± s.d. (*t*-test, \*\**P* < 0.01, *n* = 6–10). **g**,**h**, Effects of Silwet treatment on callus formation in Col-0. Heatmap of callus-forming sites (**g**) and ROS accumulation patterns by NBT and DAB staining (**h**). **i**,**j**, Callus formation in *lacs2-3*. Heatmap of callus-forming sites (**i**) and ROS staining (**j**). The number of explants forming callus at each site was visualized as a heatmap using 10 independent leaf explants (**a**, **g**, and **i**). Different letters indicate statistically significant difference (**b** and **c**). Scale bars, 0.25 cm (**a**, **d**, **f**–**j**) and 0.1 cm (**e**).

Next, we investigated how plants recognize wounding to produce ROS. Because wounding physically removes the cuticle, and cuticle defects have been shown to induce NADPH oxidase-mediated ROS production^23^, we hypothesized that cuticle removal serves as a sign of wounding. As expected, Toluidine Blue O (TBO) staining, which detects cuticle permeability, showed strong signals around the wound site in leaf explants (**Fig. 1f**). To further investigate whether cuticle defects promote ROS production relevant to callus formation, we chemically disrupted cuticle integrity by treatment of the mild surfactant Silwet prior to callus formation. Silwet-treated leaf explants exhibited callus formation and ROS accumulation throughout the entire leaf, including non-wounded regions (**Fig. 1g,h** and **Extended Data Fig. 2a**). However, ROS accumulation was markedly reduced by DPI treatment (**Extended Data Fig. 2b,c**). Similarly, the strong cuticle-defective mutant *lacs2-3* showed widespread callus formation and ROS accumulation, both of which were suppressed by DPI treatment (**Fig. 1i,j** and **Extended Data Fig. 2d-h**). In contrast, *lacs1* mutants, which exhibit a relatively mild cuticle-defective phenotype^24^, did not show altered patterns of callus formation and ROS accumulation (**Extended Data Fig. 3a-c**). These results indicate that cuticle defects serve as a sign for wound-induced ROS production, which in turn triggers the initiation of callus formation in leaf explants.

### FER restricts site of callus formation

We next examined how ROS accumulation is spatially restricted to the wound site. To identify regulators that confine ROS distribution, we screened mutants defective in RLKs containing the extracellular malectin-like domain, as these proteins function as cell wall sensors in response to mechanical stresses^25^. Among the *fer-4, herk1*, and *the1-4* mutants tested, only excised *FER*-deficient *fer-4* leaves exhibited non-site-specific callus formation (**Fig. 2a**). Consistent with this phenotype, dispersed ROS accumulation throughout the entire leaf was observed in *fer-4* (**Fig. 2b** and **Extended Data Fig. 4a,b**). Since *fer-4* mutants exhibit pavement cell bursting and ROS accumulation under high cell wall tension conditions (0.7% agar)^26^, we reduced cell wall tension by increasing agar concentration. Nonetheless, widespread ROS accumulation was still observed in *fer-4* during callus formation (**Fig. 2c** and **Extended Data Fig. 4c**), suggesting that the dispersion of ROS in *fer-4* results from misregulation of ROS distribution rather than pavement cell bursting. DPI treatment suppressed ROS accumulation, callus formation, and expression of callus formation-related genes in the wound-distal region of *fer-4* explants (**Fig. 2d-f** and **Extended Data Fig. 4d**), indicating that NADPH oxidases are the source of ROS accumulated in *fer-4*. We next examined FER protein levels and found that they were higher in wound-proximal regions than wound-distal regions (**Extended Data Fig. 4e**). However, we were unable to detect FER autophosphorylation in leaf explants during callus formation (**Extended Data Fig. 4f**). Notably, complementation of *fer-4* with a kinase-dead *FER* (*FER^K565R^*) restored the misregulated callus phenotype (**Fig. 2g**), whereas complementation with *FER* lacking the malectin-like domain (*FERΔMALA*) failed to rescue the phenotype (**Fig. 2h**). Given that FER interacts with the cell wall components via its MALA domain^27^, these findings suggest that the cell wall-interacting function of FER, rather than its kinase activity, is required for spatial regulation of ROS accumulation and restriction of callus formation to the wound site.

**Fig. 2.**
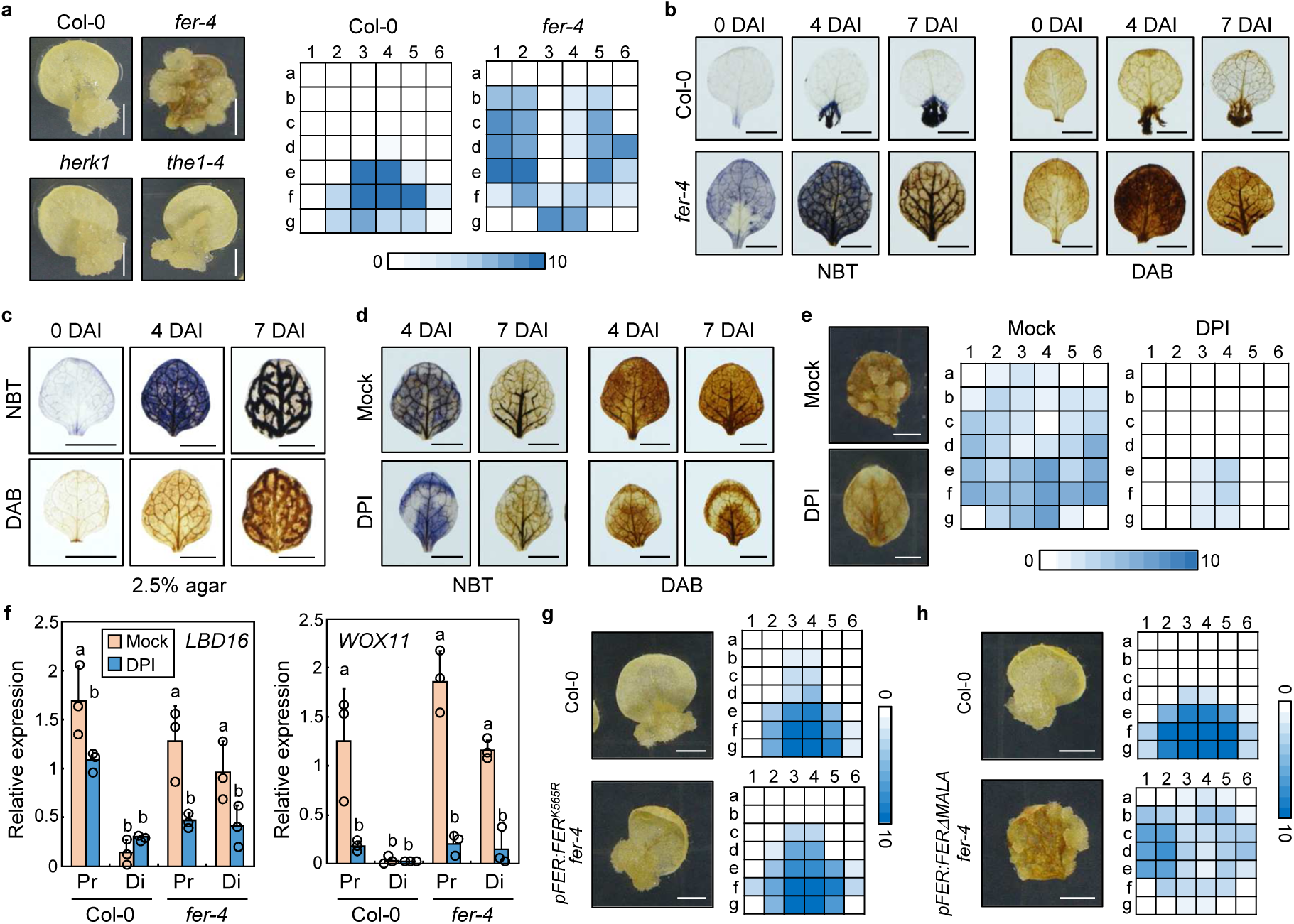
Non-site-specific ROS accumulation leads to spatially disorganized callus formation in the *fer* mutant. **a**, Screening of mutants displaying non-site-specific callus formation. Heatmaps of callus-forming sites in Col-0 and *fer-4* explants are displayed. **b**, NBT and DAB staining in Col-0 and *fer-4* explants. **c**,**d**, NBT and DAB staining during callus formation in leaf explants of *fer-4* grown on high-agar medium (**c**) or treated with DPI (**d**). **e**, Callus formation in *fer-4* explants with or without DPI treatment. Heatmap shows callus-forming sites. **f**, Expression of *LBD16* and *WOX11*. Distal (Di) and proximal (Pr) regions of leaf explants were separately harvested for gene expression analysis. Data represent mean ± s.d. from three independent biological replicates. Different letters indicate statistically significant difference (Tukey’s test, *P* < 0.05). **g**,**h**, Callus formation in *fer-4* complemented with kinase-dead or MALA domain-deleted version of FER. Leaf explants from Col-0, *pFER:FER^K565R^-GFP fer-4* (**g**), or *pFER:FERΔMALA-MYC fer-4* (**h**) seedlings were used. Heatmap shows callus-forming sites. The number of explants forming callus at each site was visualized as a heatmap using 10 independent leaf explants (**a**, **e**, **g**, and **h**). Scale bars, 0.25 cm (**a–e**, **g**, and **h**).

### FER is crucial for the development of the cuticle in leaves

Since callus formation patterns in *fer-4* closely resembled those observed in cuticle-defective explants, we hypothesized that FER is involved in cuticle formation. To verify this hypothesis, we evaluated changes in cuticle permeability between *fer-4* and the wild type using TBO staining. As a result, cuticle permeability was markedly increased in the aerial organs of 12-day-old *fer-4* compared to the wild type, and this phenotype was rescued in two complementation lines (*pFER:FER-GFP fer-4, pFER:FER-MYC fer-4*) (**Fig. 3a,b**). Next, we examined the content and composition of cutin and cuticular waxes in the aerial organs of 12-day-old plants from four lines. Total cutin monomers were reduced by approximately 25% in *fer-4* compared to the wild type, primarily due to a specific and pronounced decrease in the levels of ω-hydroxy fatty acids and 1, ω-dicarboxylic acids (**Fig. 3c** and **Extended Data Fig. 5a**). Total wax content also decreased by approximately 20% in *fer-4* relative to the wild type, with notable reductions in alkanes, aldehydes, and primary alcohols except VLCFAs (**Fig. 3d** and **Extended Data Fig. 5b**). This observation prompted us to investigate whether changes in cutin and wax loads were associated with alterations in the transcript levels of cuticle biosynthesis-related genes. Gene expression analysis showed a significant decrease in the expression of *CYP86A4, GPAT6, GPAT8, DCR, CER1, CER4*, and *SOH1* (**Fig.3e** and **Extended Data Fig. 5c**), indicating that the reduced amounts of cutin and cuticular waxes were derived from the downregulation of cuticle biosynthesis-related genes.

**Fig. 3.**
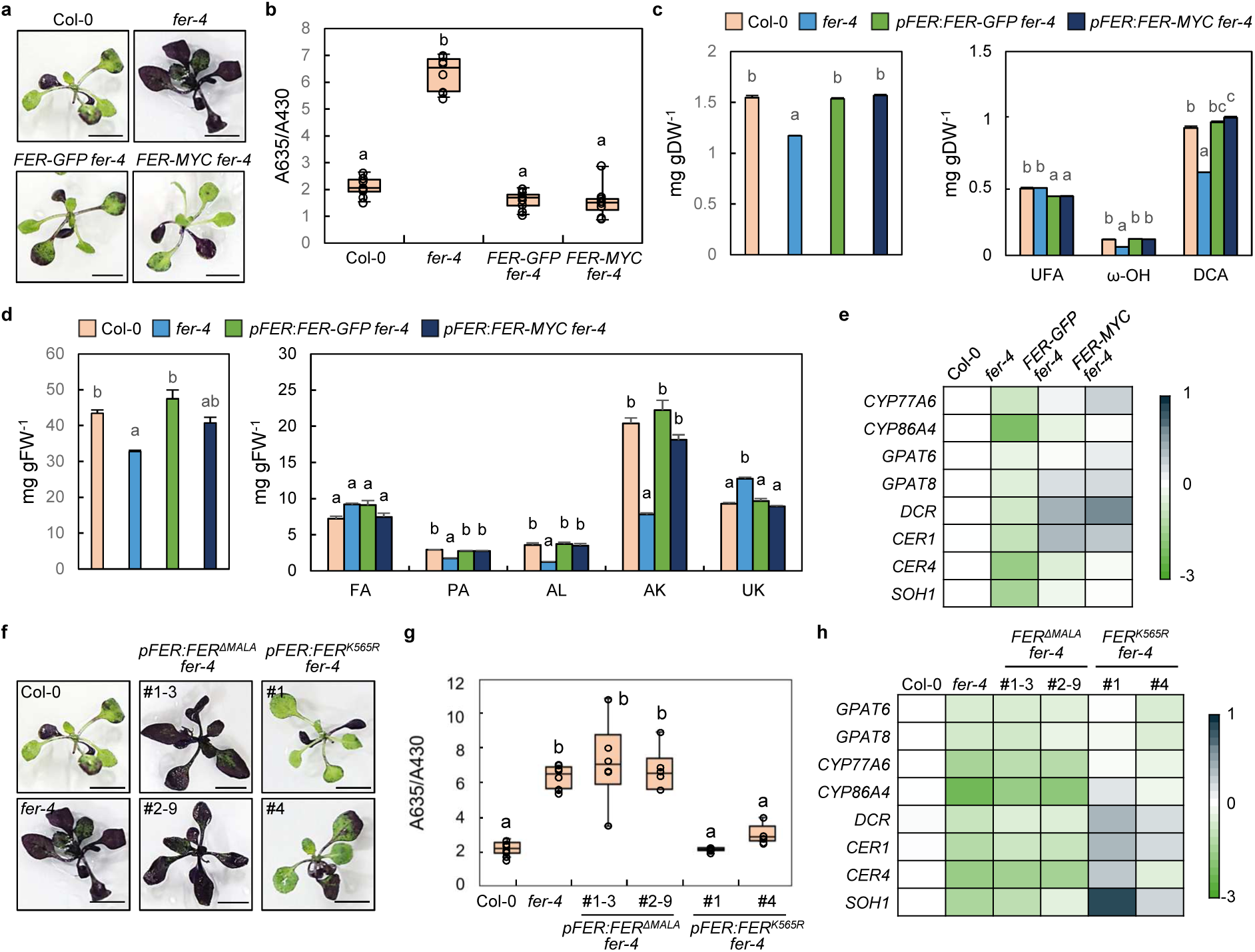
FER mediates cuticle biogenesis in epidermal cells of aerial organs. **a**,**b**, TBO staining. Aerial parts of 12-day-old plants were subjected to staining (**a**). Amount of TBO stained in each genotype was quantified and normalized by the amount of chlorophyll in eight independent biological replicates (**b**). **c**,**d**, Measurement of cutin and wax content and composition. Cutin (**c**) and wax (**d**) content of five independent biological replicates were analyzed by GC-FID. Contents of total (left panel) and individual components (right panel) are displayed. UFA, unsubstituted fatty acids; ω-OH, ω-hydroxyacids; DCA, dicarboxylic acids; AK, alkanes; AL, aldehydes; FA, fatty acids; PA, primary alcohols; UK, unknown. **e**, Expression of cuticle biosynthesis-related genes. Average relative expression was visualized as a heatmap. Scale bar, log2 fold change. **f**,**g**, TBO staining. Aerial parts of shoots were subjected to staining (**f**). Amount of TBO stained in each genotype was quantified and normalized by the amount of chlorophyll in five to eight independent biological replicates (**g**). **h**, Expression of cuticle biosynthesis-related genes. Average relative expression was visualized as a heatmap. Scale bar, log2 fold change. Aerial parts of Col-0, *fer-4*, *pFER:FER-GFP fer-4*, and *pFER:FER-MYC fer-4* seedlings were analyzed (**a**–**e**). Aerial parts of Col-0, *fer-4*, *pFER:FERΔMALA-MYC fer-4* and *pFER:FER^K565R^-GFP fer-4* were analyzed (**f**–**h**). For **c** and **d**, data represent mean ± s.e. For **b**–**d** and **g**, different letters indicate statistically significant difference (Tukey’s test, *P* < 0.01). Scale bars, 0.5 cm (**a** and **f**).

To determine whether the MALA domain or the kinase domain of FER is involved in cuticle formation, we performed TBO staining of *pFER:FER^K565R^ fer-4* and *pFER:FERΔMALA fer-4* lines. Similar to callus formation, only the complementation of *fer-4* with a MALA-deleted FER failed to rescue cuticle defects (**Fig. 3f,g**). Genes that were downregulated in *fer-4* compared to the wild type were also repressed in *pFER:FERΔMALA fer-4* but they were partly rescued or even upregulated in *pFER:FER^K565R^ fer-4* (**Fig. 3h**). This demonstrates that the MALA domain is critically involved in cuticle formation, likely independent from the kinase activity of FER. This result also aligns with the above results, where only the MALA domain-deleted FER caused unorganized callus formation (**Fig. 2g,h**).

### FER mediates wound-induced cuticle development in mesophyll cells

Our observations that wound-produced ROS did not propagate beyond the BC suggested the presence of barriers that confine ROS near the wound site (**Fig. 1e**). Based on the widespread ROS accumulation observed in cuticle-defective mutants (**Fig. 1j** and **Fig. 2b**) and a previous report showing that cuticular waxes accumulate in response to wounding in leaves^9^, we hypothesized that this barrier is the cuticle. Supporting this hypothesis, the expression of cuticle biosynthesis genes was specifically upregulated in wound-proximal regions but markedly suppressed in *fer-4* during callus formation (**Fig. 4a**). To pinpoint the site of FER-mediated, wound-responsive cuticle development, we examined the deposition site of demethylesterified pectin, which interacts with the MALA domain of FER for its activation^27^. Ruthenium red staining revealed that pectin demethylation occurred at the BC (**Fig. 4b** and **Extended Data Fig. 6a**), which coincides with the site where ROS propagation is halted (**Fig. 1e**). Moreover, the expression of *PME2* and *PME3*, which encode pectin methylesterases that induce pectin demethylesterification, was significantly upregulated in wound-proximal regions during callus formation (**Fig. 4c**), supporting the occurrence of pectin demethylation at the BC.

**Fig. 4.**
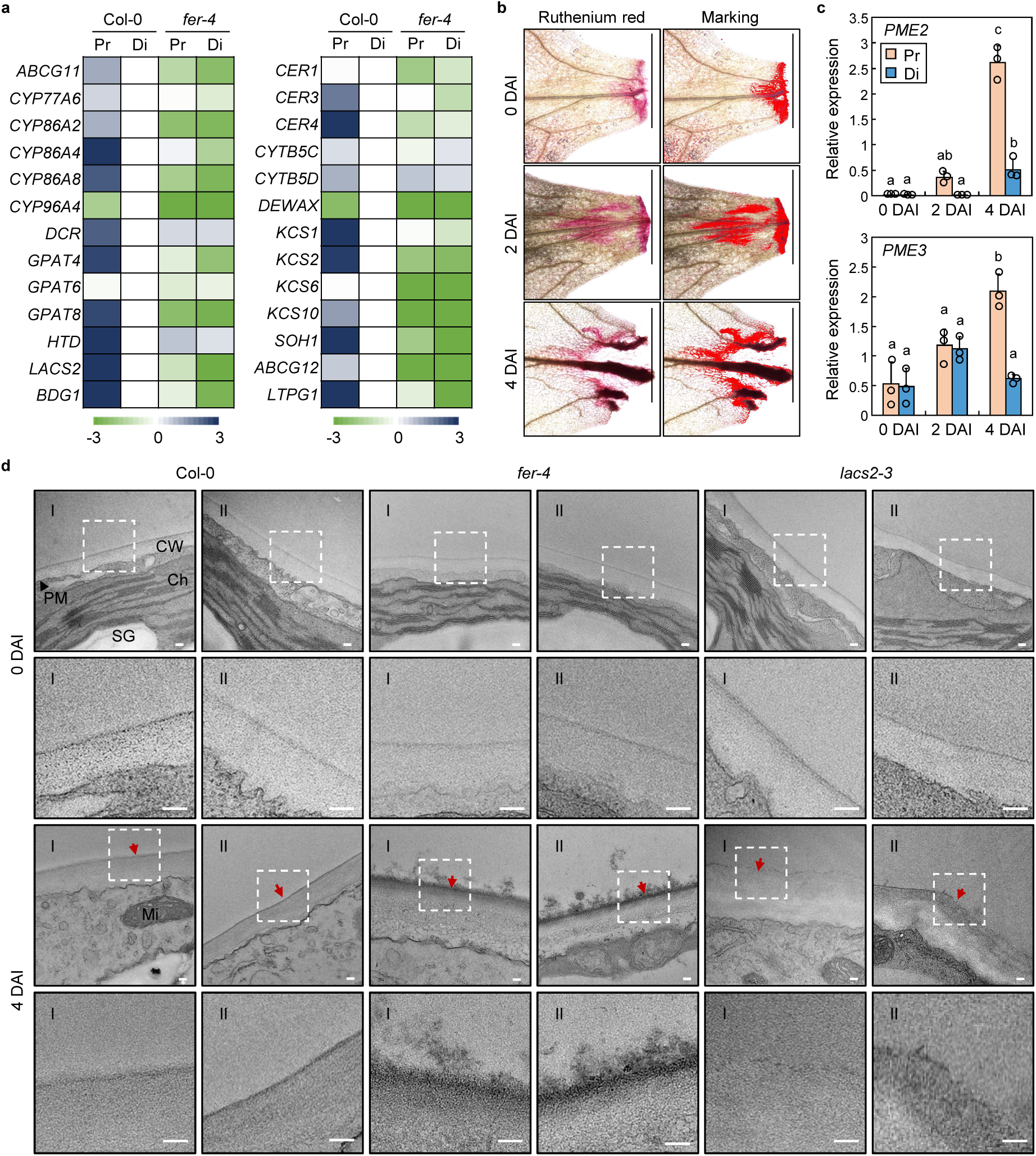
FER induces cuticle formation in mesophyll cells near the wound site. **a**, Expression of genes involved in cuticle biosynthesis in leaf explants during callus formation. Pr and Di regions were analyzed separately. Average expression values of cutin (left panel) and wax (right panel) biosynthesis-related genes from three independent biological replicates are displayed as a heatmap, relative to Col-0 Di. Scale bars, log2 fold change. **b**, Ruthenium red staining. Col-0 leaf explants were subjected to staining during callus formation. Stained regions were highlighted using ImageJ software (Marking). **c**, Expression of *PME*s. Pr and Di regions of Col-0 leaf explants were separately analyzed. Data represent mean ± s.d. from three independent biological replicates. Different letters indicate statistically significant difference (Tukey’s test, *P* < 0.05). **d**, TEM imaging of mesophyll cells near the wound site. Mesophyll cells in BC or the corresponding regions near the wound site of CIM-incubated leaf explants were subjected to TEM imaging. Cells in direct proximity to the wound site (I) and peripheral cells located slightly distal to the wound (II) were analyzed. CW, cell wall; PM, plasma membrane; Ch, chloroplast; SG, starch granule; Mi, mitochondria. Lower panels for each DAI show enlarged views of the dotted areas. Red arrows indicate the cuticle. Scale bars, 0.1 cm (**b**) and 100 nm (**d**).

The cuticle is an epidermal cell-specific structure in the aerial part of plants, so we thought that it must be formed in the mesophyll cells to function as a barrier to restrict further ROS propagation. To investigate whether a cuticle is formed at the BC during callus formation, we sectioned wound-proximal regions (**Extended Data Fig. 6b,c**) and analyzed the BC in Col-0, as well as the corresponding areas in *fer-4* and *lacs2-3*, using transmission electron microscopy (TEM). At 0 DAI, there were no observable differences among Col-0, *fer-4*, and *lacs2-3*, where only an unidentified faint thin layer was observed on the surface of mesophyll cells. However, at 4 DAI, a significant difference became evident. A compact and vivid layer, presumed to be a cuticle, reinforced on the mesophyll surface of Col-0 (**Fig. 4d**). In contrast, the layer in *fer-4* was disrupted, appearing irregular and swollen, some parts detaching apart. In *lacs2-3*, the layer thickness seemed similar to the wild type, but displayed a discontinuous and patchy phenotype. These results suggest that a cuticle is newly formed on the surface of mesophyll cells at the BC, indicating transdifferentiation of these cells into epidermal cells.

### PCD ensures ROS confinement near the wound site

To gain further insight into the mechanism that confines callus formation to the wound region, we examined morphological changes in leaf explants during callus formation. Notably, site-specific programmed cell death (PCD) occurred along mesophyll cells within the region of callus formation (RC) (**Fig. 5a**). This PCD was accompanied by cell detachment, a hallmark of apoptosis^28^. Because apoptotic cell death can be triggered by hydroxyl radicals^29^, we stained hydroxyl radicals using Rhodamine B hydrazide (RBH) and found that they accumulated specifically in the RC (**Extended Data Fig. 7a**). To further investigate the role of hydroxyl radicals in PCD and callus formation, we treated explants with N,N’-dimethylthiourea (DMTU), a hydroxyl radical scavenger^30^ (**Extended Data Fig. 7b**). DMTU treatment suppressed PCD in RC and caused leakage of ROS into wound-distal regions (**Fig. 5a** and **Extended Data Fig. 7c,d**). Consistent with the expanded ROS distribution, the area of callus formation also extended into wound-distal regions (**Fig. 5b**). Co-treatment with DPI and DMTU suppressed ROS accumulation (**Fig. 5c** and **Extended Data Fig. 7e**), indicating that the dispersed ROS is originated from NADPH oxidases.

**Fig. 5.**
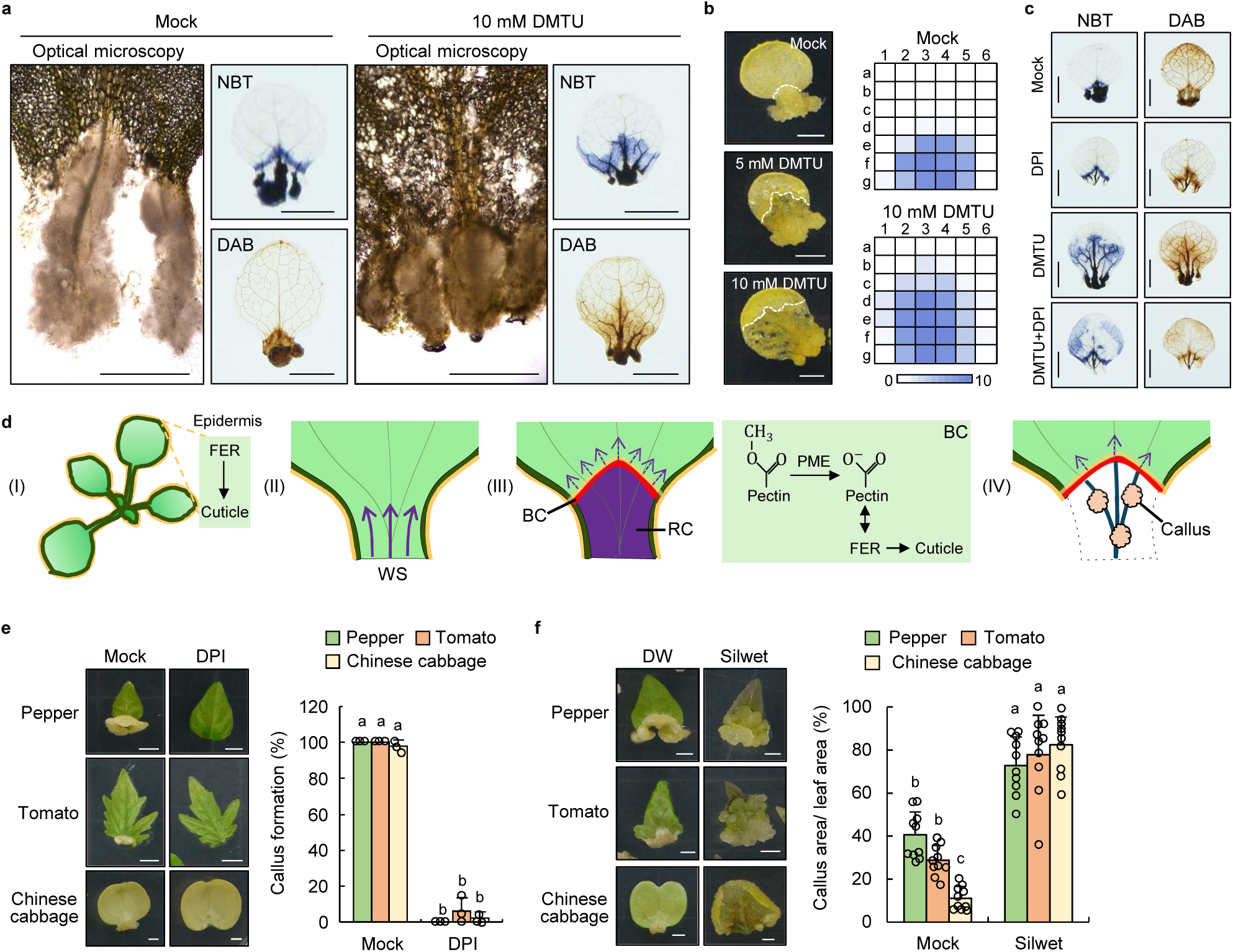
PCD ensures the restriction of ROS propagation to the distal part. **a**, Morphology and ROS accumulation during callus formation with DMTU treatment. Col-0 leaf explants at 7 DAI were analyzed. Scale bars, 0.1 cm (optical microscopy); 0.25 cm (NBT and DAB staining). **b**, Effects of DMTU on callus formation in Col-0. Heatmap shows callus-forming sites. The number of explants forming callus at each site was visualized as a heatmap using 10 independent leaf explants. **c**, Effects of DPI on ROS accumulation in DMTU-treated explants. Col-0 leaf explants at 7 DAI were subjected to NBT and DAB staining. **d**, Model for cuticle-defined, site-specific callus formation. Arrows indicate ROS propagation and the yellow lines indicate cuticle layer. (I) Developing leaves before wounding. (II) Leaf explants immediately after wounding. (III) 2 to 4 DAI during CIM incubation. (IV) Callus formation at VT after 4 DAI. WS, wound site. **e**, Effects of DPI on callus formation in crop species. Leaf explants of pepper, tomato, and Chinese cabbage at 11, 8, and 11 DAI, respectively, are displayed. Percentage of explants forming callus is displayed. Data represent mean ± s.d. (*n* = 3). **f**, Effects of Silwet on leaf explants in crop species. Leaf explants of pepper, tomato, and Chinese cabbage at 18, 14, and 11 DAI, respectively, are displayed. Percentage of callus area relative to leaf area is displayed. Data represent mean ± s.d. (*n* = 10). For **e** and **f**, different letters indicate statistically significant difference (Tukey’s test, *P* < 0.05). Scale bars, 0.25 cm (**b**, **c**, **e**, and **f**).

Based on our findings, we present a model summarizing this mechanism (**Fig. 5d**). FER facilitates the development of the cuticle in leaves, possibly triggered by the exposure of their epidermal cells to air and the mechanical stress generated from cell expansion. Upon wounding, the cuticle is disrupted, leading to generation of ROS at the wound site. These ROS initially propagate toward distal regions. During this process, wound-induced pectin demethyl-esterification in the cell wall activates the plasma membrane-localized FER, which in turn promotes *de novo* cuticle formation at the BC. The newly formed cuticle acts as a primary barrier that limits ROS propagation. Subsequently, locally confined ROS induce apoptotic PCD in the surrounding mesophyll cells, physically separating them from adjacent tissues and thereby preventing further propagation of ROS to distal regions. Within the remaining vasculature, which would be resistant to PCD, ROS persist and serve as a cue to initiate callus formation.

To evaluate whether this spatial regulation mechanism is conserved in crops, we examined callus formation in tomato, pepper, and Chinese cabbage. In all tested species, callus formation was initiated at the wound site and was markedly suppressed by DPI treatment (**Fig. 5e**). In addition, treatment with Silwet resulted in widespread callus formation (**Fig. 5f**), indicating that the cuticle-ROS regulatory axis restricting callus formation to wound sites is broadly conserved among dicot crops.

## Discussion

The plant cuticle is a hydrophobic barrier that imparts impermeability to water, gases, and various harmful substances^2^. This property was pivotal for the terrestrialization of early land plants, enabling survival under fluctuating humidity and temperature and remarkably elevated oxygen conditions after transitioning from aquatic environments^4^. In land plants, the cuticle remains a dynamic interface with the environment, and its composition and structure are rapidly modulated in response to diverse abiotic stresses such as drought, salinity, temperature extremes, and UV-B radiation^31–33^. In particular, alkanes, a major component of cuticular waxes, was shown to be negatively correlated with cuticle permeability and was profoundly induced upon wounding or drought conditions^10,31,34^. These findings suggest that cuticle remodeling plays an essential role in reinforcing epidermal cell barrier in response to external signals. In this study, for the first time to our knowledge, we reveal that the barrier function of cuticle also contributes to restricting further propagation of ROS.

Extensive studies over the past decades have revealed finely tuned pathway of embryonic cuticle surveillance, involving certain RLKs. During seed maturation, GSO1/2 with coreceptor SOMATIC EMBRYOGENESIS RECEPTOR-LIKE KINASEs perceive TWISTED SEED1 (TWS1) peptide, which is activated by endosperm-specific ALE1 subtilase. TWS1-mediated embryonic cuticle reinforcement continues until the cuticle layer is intact enough to block TWS1 secretion from the embryo^7,35–37^. As the seed prepares germination, endosperm-derived signaling peptides penetrate the embryonic cuticle, where they are perceived by GSO1/2 and PSY1 RECEPTOR to promote cuticle development during the embryo-to-seedling transition^38^. Our study identified FER as the first RLK upstream of cuticle formation during the post-germination stage. Interestingly, cuticle development via FER depended on its MALA domain but not on its kinase activity (**Fig. 3f-h**). It was shown that FER can act as a scaffold to regulate immune signaling by inducing complex formation between FLAGELLIN-SENSING 2/EF-TU RECEPTOR and their coreceptor BRASSINOSTEROID INSENSITIVE 1-ASSOCIATED KINASE 1^39^. FER-mediated cuticle formation may also accompany FER’s scaffold function, which stabilizes proteins involved in cuticle-forming pathways. Detailed molecular mechanism of how FER regulates cuticle formation should be further investigated.

It was proposed that increased cuticle permeability induces ROS production. Cuticle biosynthesis mutants *bdg* and *lacs2-3* exhibited increased fluorescence following DCF-DA infiltration, while a cuticle defective *eca2* showed an enlarged stained area in DAB and NBT assays compared to the wild type^23,41^. These phenomena have been interpreted as the plant perceiving a compromised cuticle barrier as a signal for pathogen invasion and subsequently inducing ROS production as a defense response^42,43^. While ROS serve as important signaling molecules involved in plant immunity, their strong oxidizing nature necessitates tight regulation to prevent cellular damage as can be seen in localized ROS production around manual wounding site and restricted hypersensitive response (HR), a form of PCD, following pathogen infection^44,45^. During HR, plants recognize microbe-associated molecular patterns at the cell surface as well as injected effectors inside host cells, leading to NADPH oxidase-dependent ROS production^46^. The generated ROS accumulate only around the infection site, thereby restricting HR to infection-proximal areas^44^. A similar spatial restriction of ROS and PCD was observed in wound-proximal regions in our study. Taken together, based on previous findings that cuticle disruption can trigger plant defense-related ROS production, along with our data showing that the cuticle restricts ROS distribution, it is likely that the cuticle plays a dual role: not only as a ROS inducer upon damage but also as a physical barrier for spatial regulation of ROS in both wound-healing and immune responses.

How does cuticle block ROS propagation? When ROS are produced, the apoplastic hydrogen peroxide activates the plasma membrane-localized ROS sensor HYDROGEN PEROXIDE Ca^2+^ INCREASES 1 (HPCA1), which induces calcium influx into the cytosol. The resulting increase in cytosolic Ca^2+^ leads to activation of CALCIUM-DEPENDENT PROTEIN KINASEs and NADPH oxidases. This process stimulates apoplastic ROS propagation, leading to so-called ‘ROS wave’ across neighboring cells^46–48^. Apoplastic ROS can diffuse freely through the cell wall; however, if a cuticle layer is formed as an external barrier that restricts water and gas exchange, it likely impedes apoplastic ROS from reaching and activating HPCA1 in neighboring cells.

Living organisms have evolutionally conserved wound-healing mechanisms to minimize microbial infection and restore tissue integrity. In animals, ROS are produced in wound sites and play pivotal roles in haemostasis, immune responses, and the regeneration of cellular components^49^. Both the levels and spatial distribution of ROS are tightly regulated to focus the wound-healing process at the wound site and to prevent excessive oxidative damage^49,50^. A study using planarian flatworms further demonstrated that muscle contraction occurs around the wound region, where ROS accumulate, to minimize wound surface area^50^. Our findings suggest that plants possess conceptually similar wound-healing strategies. In Arabidopsis leaves, ROS accumulated specifically at the wound site to facilitate healing through callus formation, while a cuticle formed around the wound region to confine ROS accumulation and limit the wound-healing response to damaged area (**Fig. 5d**). Although the molecular mechanisms differ between animals and plants, both have likely evolved analogous strategies to protect intact regions while regenerating tissues in damaged areas through the controlled use of highly oxidative molecules such as ROS.

## Acknowledgements

The authors thank to prof. Sarah M Assmann for sharing *pFER:FER-GFP fer-4* and *pFER:FER^K565R^-GFP fer-4* seeds. This work was supported by grants from the National Research Foundation of Korea (RS-2021-NR058215 and RS-2022-NR070837 to M.C.S.; RS-2022-NR075618 to H.S.K.; RS-2023-NR076489 to H.J.L.), Rural Development Administration of Korea (RS-2024-00322275 to H.S.K.), and Korea Research Institute of Bioscience and Biotechnology (KGM1002521 to H.J.L.; KGM1082511 to H.S.C.).

## Author contributions

G.K., J.C., M.C.S., and H.J.L. conceived the study and wrote the manuscript. G.K. performed experiments related to callus formation. J.C. performed experiments related to cuticle development. J.C. and R.J.K. performed TEM analysis. E.O. generated the complementation lines of *fer-4*. S.Y.S. performed callus formation assays using RLK mutants. H.S.K. and H.S.C. provided scientific discussions and materials for the study.

## Competing interests

Authors declare that they have no competing interests.

## Correspondence and requests for materials

should be addressed to Mi Chung Suh or Hyo-Jun Lee.

## Methods

### Plant materials and growth conditions

The *Arabidopsis thaliana* (Arabidopsis) ecotype Columbia-0 (Col-0) was used as the wild-type control. Mutant lines including *lacs1-1* (CS65774), *lacs1-2* (CS65775), *lacs2-3* (CS65776), *fer-4* (N69044), *the1-4* (CS829966), *rbohDF* (N9558), and *herk1* (SALK_008043C) were obtained from the Nottingham Arabidopsis Stock Centre (NASC, Nottingham, United Kingdom) and the Arabidopsis Biological Resource Center (ABRC, Ohio State University, Columbus, OH, United States). The *pCYCB1;1:GUS*^51^*, pWOX11:GUS*^21^, *pFER:FER-MYC fer-4*^52^, *pFER:FER^K565R^-GFP fer-4*^53^, and *pFER:FER-GFP fer-4*^53^ were reported previously. To generate *pFER:FERΔMALA-MYC fer-4*, the *FER* gene in pFER:FER-MYC vector^52^ was substituted with MALA-deleted *FER* (deletion of 103 to 570 bps) using Q5 Site-Directed Mutagenesis kit (New England Biolabs, cat. no. E0554S) and transformed into *fer-4* using *Agrobacterium* floral dip method^54^.

For callus formation, Arabidopsis, Chinese cabbage (*Brassica rapa* cv. Jangsaeng-3-ho), tomato (*Solanum lycopersicum* cv. Super Doterang), and pepper (*Capsicum annuum* cv. Nokkwang) were used in this study. Seeds were germinated and grown on half-strength Murashige and Skoog (1/2 MS) medium supplemented with 0.68% (w/v) plant agar. Seeds of Arabidopsis and Chinese cabbage were stratified at 4□°C in darkness for 3 days prior to cultivation. After stratification, seedlings were grown under long-day conditions (16 hours light and 8 hours dark) for 12 days (Arabidopsis) or 5 days (Chinese cabbage). Tomato and pepper seeds were incubated in the dark for 4 days and 7 days, respectively, followed by cultivation under long-day conditions for 8 days (tomato) or 7 days (pepper). All seedlings were grown in growth room at 24□°C with 40-50% humidity.

For cuticle analysis, TBO staining and quantitative reverse transcription PCR (RT-qPCR) of cuticle-related genes in 12-day-old plants, plants were treated as follows. Seeds were sterilized with 75% ethanol + 0.05% Triton X-100 for 30 seconds, washed with 100% ethanol and completely dried on filter paper before sowing on 0.6% 1/2 Murashige and Skoog media containing 1% sucrose. Plants were stratified in 4 □ for 3 days and transferred to a finely controlled growth room kept of 23 □, 50-60% humidity and 16 h light/8 h dark.

### Callus induction

Callus induction was performed in Arabidopsis, Chinese cabbage, tomato, and pepper. Leaf explants were prepared by excising the first pair of true leaves (Arabidopsis, tomato, pepper) or cotyledons (Chinese cabbage) at the petiole–blade junction and placing them adaxial side down on callus induction medium (CIM), which consisted of B5 basal salts, 3% (w/v) sucrose, 0.05□mg□L^-1^ kinetin, 0.5□mg□L^-1^ 2,4-dichlorophenoxyacetic acid (2,4-D), and 0.68% (w/v) plant agar, adjusted to pH 5.7 with NaOH. Explants were incubated in darkness at 24□°C for species-specific durations: 14 days for Arabidopsis, 11 days for Chinese cabbage, 8 days for tomato, and 11 days for pepper. In Arabidopsis, additional experimental conditions were applied in several experiments. Leaf explants were harvested at different time points depending on the purpose of the analysis, as indicated in the figure legends.

### Generation of heatmap

To visualize spatial patterns of callus formation on leaf explants, images of individual explants were taken under uniform orientation and light conditions. Each image was divided into a 6 × 7 grid, and each section was manually scored for the presence or absence of callus. Ten explants were analyzed, and the number of explants showing callus in each section was counted. The frequency and spatial distribution of callus formation were visualized as a heatmap using Microsoft Excel. To visualize gene expression as a heatmap, transcript levels of genes were measured by RT-qPCR and log2-transformed relative average expression levels were used. Microsoft Excel was used to construct the heatmap.

### Pharmacological treatment

Pharmacological treatments were applied to leaf explants during callus formation. Three compounds were used: DPI (Sigma-Aldrich, cat. no. D2926), DMTU (Sigma-Aldrich, cat. no. D188700), and Silwet L-77 (PlantMedia, cat. no. 30630216-1). Mock explants were treated with the same solvent used for each compound, without the active chemical. For DPI treatment, leaf explants were cultured on CIM containing 1□μM DPI in Arabidopsis, 20□μM in Chinese cabbage, and 10□μM in tomato and pepper. Incubation durations were 14 days for Arabidopsis, 11 days for Chinese cabbage and pepper, and 8 days for tomato. For Silwet L-77 treatment, explants were placed on CIM and then sprayed with species-specific concentrations: 0.02% for Arabidopsis, 0.08% for Chinese cabbage and pepper, and 0.04% for tomato. The explants were incubated for 14 days (Arabidopsis), 11 days (Chinese cabbage), 8 days (tomato), or 18 days (pepper). For DMTU treatment, explants were cultured on CIM supplemented with 10□mM DMTU. This treatment was primarily applied to Arabidopsis. Time point-specific analyses including phenotypic evaluation, ROS distribution, and gene expression profiling were performed depending on the experimental objective, as indicated in the figure legends.

### GUS staining

GUS staining was performed using *pWOX11:GUS* and *pCYCB1;1:GUS* transgenic lines. The first true leaves were excised and cultured on CIM containing 1□μM DPI for 4 days prior to staining. Explants were harvested and fixed in 90% cold acetone on ice for 20 minutes and rinsed twice with rinse solution containing 50□mM sodium phosphate (pH 7.2), 0.5 mM K_3_Fe(CN)_6_, and 0.5 mM K_4_Fe(CN)_6_. Tissues were then incubated in staining solution [rinse solution additionally containing 1□mM X-Gluc (Duchefa), 10 mM EDTA, 0.1% Triton X-100] at 37□°C overnight in darkness. After staining, explants were immersed in 70% ethanol for 24 hours to remove chlorophyll. Images were produced using a Nikon D750 digital camera.

### TBO staining

Cuticle permeability of leaf explants was evaluated by TBO (Sigma-Aldrich, cat. no. T3260) staining. Explants cultured on CIM were harvested at the time points indicated in the figure legends and immersed in 0.02% (w/v) TBO solution for 5 minutes at room temperature. After staining, samples were gently rinsed with distilled water and immediately photographed using a Nikon D750 digital camera. Signal intensity was quantified using ImageJ software.

For seedlings, TBO staining and quantification were conducted following methods in Li et al. (2016)^55^. Aerial parts of 12-day-old plants were excised and immersed in 0.05% TBO (w/ 0.01% Tween 20) for indicated amount of time, washed in DW, and photographed under digital camera. Sequentially, for TBO quantification, each replicate of plant samples was ground with extraction buffer (200 mM Tris-HCl pH 7.5, 250 mM NaCl, 25 mM EDTA) and 2 volume EtOH was added. After centrifugation at 13,000 rpm for 1 min, the supernatant was transferred and the A635/A430 ratio was measured using spectrophotometer to quantify the amount of TBO, normalized by the amount of chlorophyll.

### NBT and DAB staining

ROS staining was performed on leaf explants cultured on CIM under various conditions, including different genotypes, time points, and pharmacological treatments, as detailed in the figure legends. NBT (Thermo Scientific, cat. no. B23792.88) staining was performed to detect superoxide anion and DAB (Sigma-Aldrich, cat. no. D8001) staining was performed to detect hydrogen peroxide. For DAB staining, leaf explants were immersed in a freshly prepared solution containing 40□mg DAB and 20□μL Tween 20 in 40□mL distilled water (pH adjusted to 3.0 with HCl). Explants were vacuum-infiltrated for 2 minutes and incubated in the dark at 24□°C for approximately 6 hours with gentle shaking. Stained tissues were then cleared of chlorophyll in 70% ethanol. For NBT staining, explants were immersed in a solution containing 70□mg NBT and 13□mg sodium azide in 20□mL of 10□mM potassium phosphate buffer (pH 7.8). Samples were vacuum-infiltrated for 2 minutes and incubated in the dark at 24□°C for 2 hours with shaking. After staining, tissues were cleared of chlorophyll in 70% ethanol. Images were acquired using a Nikon D750 digital camera, and signal intensity was quantified using ImageJ software.

### Ruthenium red staining

To visualize de-methyl esterified pectin in leaf explants, leaf explants were incubated on CIM for the time periods indicated in the figure legends. Explants were cleared of chlorophyll using acid alcohol solution (glacial acetic acid:95% ethanol = 1:3, v/v) until visibly decolorized, then rehydrated through a graded ethanol series (70% - 60% - 50%) to minimize osmotic stress and preserve tissue structure. Rehydrated tissues were incubated in 0.01% (w/v) Ruthenium Red (Sigma-Aldrich, cat. no. R2751) solution (prepared in distilled water) for 10 minutes at room temperature and subsequently rinsed with distilled water. Stained samples were immediately examined under an optical microscope to assess the localization of de-methyl esterified pectin.

### Analysis of cutin and wax content

For cuticular wax analysis, the aerial parts from 12-day-old plants were excised and gently shaken in 2 mL chloroform for 30 seconds. Residual wax was collected with an additional 500 μl of chloroform. Heneicosanoic acid, 1-tricosanol, and octacosane were added as internal standards. Dried residues under a stream of nitrogen gas were redissolved in 100 μl pyridine (Thermo Fisher Scientific) and 100 μl N,O-bis(trimethylsilyl)trifluoroacetamide (BSTFA; Sigma-Aldrich) and incubated at 120 °C in a sand bath for 30 minutes for derivatization. Solvents were evaporated again under nitrogen gas, and the residues were dissolved in a 1:1 (v/v) mixture of heptane and toluene. Samples were analyzed by gas chromatography (GC-2010; Shimadzu) equipped with a DB-5 column (60 m × 0.32 mm i.d., 0.25 μm film thickness; Agilent) as described in Go et al. (2014)^56^. For cutin polyester analysis, wax-extracted plant samples were boiled in 80 □ isopropanol for 10 minutes and incubated in isopropanol on shaking incubator until all chlorophylls were removed. Afterwards, plants were ground using mortars and pestles and delipidated in gradual exchange of solvents containing chloroform and methanol (2:1, 1:2). Plant residues were finally washed in methanol and completely dried overnight in a vacuum. 0.01% BHT was added to all solvents for delipidation steps. Dried residues were mixed with methyl heptadecanoate and ω-pentadecalactone as internal standards, and incubated with reaction solvents containing methanol, 28% sodium methoxide and methyl acetate for 2 hours at 60 □. Lipids were extracted in dichloromethane, washed with 0.9% NaCl (in 100 mM Tris-HCl pH 8.0) and sodium sulfate anhydrous, concentrated under nitrogen gas and finally dissolved in heptane/toluene (1:1, v:v) to be analyzed using GC-FID as described in Kim et al. (2025)^57^. Individual wax and cutin compounds were identified through comparison with internal standards.

### RBH staining

To detect hydroxyl radicals in leaf explants during callus induction, explants cultured on CIM for 3 days were harvested and rinsed twice with 1/2 MS liquid medium. The explants were then incubated in 50□μM RBH (TCI, cat. no. R0260) solution for 30 minutes under dark conditions with gentle shaking. After staining, the samples were washed twice with 1/2 MS liquid medium and immediately imaged using an LSM 800 confocal microscope (Carl Zeiss). For RBH detection, excitation and emission wavelengths were set to 543□nm and 565□nm, respectively.

### RT-qPCR

For gene expression analysis under callus induction conditions, CIM-cultivated leaf explants were collected at various time points either as whole tissues or after dissection into proximal (the region containing the wound site) and distal (the region distant from the wound site) regions prior to RNA extraction. Total RNA of leaf explants was extracted using TRIzol reagent (Thermo Fisher Scientific) according to the manufacturer’s instructions. Complementary DNA (cDNA) was synthesized from 1□μg of total RNA using the AccuPower® CycleScript RT PreMix (Bioneer, cat. no. K-2044). Quantitative PCR was carried out with the TOPreal SYBR Green qPCR PreMIX (Enzynomics, cat. no. RT500M) on a CFX Duet Real-Time PCR System (Bio-Rad, 12016265). Primer sequences used in this study are provided in **Supplementary Table 1**. Gene expression levels were normalized to *UBQ10*, which served as the internal reference.

For genes related to cuticle biogenesis, Twelve-day-old plants were finely ground with liquid nitrogen and the RNA was extracted with XENOPURE^TM^ Total RNA Purification Kit following manufacturer’s instructions. cDNA was synthesized with 2 μg RNA as template using GoScript^TM^ Reverse Transcriptase (Promega) following manufacturer’s instructions. Quantitative PCR was proceeded with CFX Opus 96 real time quantitative PCR system (Bio-rad). *PP2AA3* (At1g13320) was used to evaluate RNA quantity and quality and to serve as a normalization gene.

### Imaging of sectioned area

Leaf explants cultured on CIM for 2 days were stained with NBT and used for paraffin sectioning. Samples were dehydrated through a graded ethanol series (70%, 80%, 85%, 90%, 95% and 100%; 1 h each step, 100% ethanol repeated three times), cleared in Histo-Clear (National Diagnostics, cat. no. NAT1334), and embedded in Paraplast (Leica, Wetzlar,Germany, cat. 39601006). Paraffin blocks were sectioned at a thickness of 10□μm using an Epredia HM 340E Electronic Rotary Microtome (Thermo Fisher Scientific, Epredia). Sections were deparaffinized with Histo-Clear, rehydrated through a graded ethanol series (1:1 Histo-Clear:ethanol solution, 100%, 95%, 70%, and 50% ethanol, and distilled water) and mounted using Permount mounting medium (Fisher chem., cat. SP15-100). Slides were scanned using a slide scanner (Carl Zeiss, Axio Scan Z1).

### Immunoprecipitation and immunoblot analysis

Leaf explants of *pFER:FER-MYC* transgenic Arabidopsis cultured on CIM for 14 days were dissected into proximal and distal regions, ground in liquid nitrogen, and lysed in IP buffer (100□mM Tris-HCl pH 7.5, 75□mM NaCl, 10% glycerol, 1□mM EDTA, 0.1% Triton X-100, and protease inhibitors). Lysates were cleared by centrifugation and the supernatant was used. For immunoprecipitation, anti-MYC antibody (Millipore, cat. no. 05-724) was pre-bound to Protein G Magnetic Beads (Bio-Rad, cat. no. 1614023) and incubated with the extracts at 4□°C. The beads were washed with wash buffer (IP buffer without protease inhibitors). After washing, bound proteins were eluted in SDS sample buffer [150 mM Tris-Cl, pH 6.8, 4.8% (w/v) SDS, trace amout of bromophenol blue, 24% (v/v) glycerol, and 672 mM β-mercaptoethanol].

For immunoblot assays, proteins were resolved by SDS-PAGE, transferred to PVDF membranes (Millipore), and immunoblotted with antibodies against phosphoserine (Abcam, cat. no. ab9332), phosphotyrosine (Abcam, cat. no. ab179530), phosphothreonine (Abcam, cat. no. ab218195), and MYC. For chemiluminoscence detection of proteins, membranes were blotted using HRP-conjugated secondary antibodies (Abcam), and then subjected to treatment of ECL reagents (BioD, cat. no. A0031). Molecular Imager® Gel Doc™ XR+ System (Bio-Rad, cat. no. 1708195EDU) was used to analyze immunoblots.

### TEM analysis

Leaf explants of Col-0, *fer-4*, and *lacs2-3* cultured on CIM for 0 or 4 days were carefully harvested and fixed overnight at 4°C in the dark using a fixation solution containing 2% paraformaldehyde, 2.5% glutaraldehyde, and 0.1 M sodium cacodylate buffer (pH 7.4). Samples were subsequently washed 5–6 times with 0.1□M sodium cacodylate buffer (pH 7.4) for 10 minutes each. Post-fixation was performed using 1% osmium tetroxide (OsO_4_) at 4°C for 1 hour to enhance membrane contrast and stabilize lipid components. The samples were then washed again 7–10 times with 0.1□M sodium cacodylate buffer. After fixation, samples were dehydrated through a graded ethanol series (10%, 20%, 30%, 40%, 50%, 60%, 70%, 80%, 90% and 100%; 30 minutes each step, 100% ethanol repeated three times) and incubated overnight at 4°C in the dark with gentle inverting. Dehydrated samples were gradually embedded with Spurr’s epoxy resin (medium hardness, Ted Pella) with sequential exchange of EtOH:resin=2:1, 1:1, 1:2 (each 2 hours incubation), and finally 100% resin o/n. Samples were hardened for 36∼48 hours at 60 □. Resin-embedded samples were sectioned into 80–100 nm slices using an ultramicrotome (RMC Products) and then subjected to light microscope (For Extended Data Fig. 6c) and TEM imaging. For light microscope imaging, the floating sections were carefully mounted onto slide glasses, fixed by water-evaporation on heat block, and then 0.05% TBO solution was evenly applied to the surface of each slide. The TBO solution was completely dried on a heat block. Nonspecific dye was washed off with DW and TBO-stained tissues were observed and imaged under a light microscope with integrated camera (Leica ICC50 HD). For TEM imaging, sections were placed on a grid and stained with uranyl acetate and lead citrate prior to observation with TEM (Jeol, JEM-2100F).

### Optical microscopy

Bright-field microscopy was performed using an upright microscope (BX53, Olympus) equipped with a camera adaptor (U-CMAD3, Olympus). Tissue morphology near the wound site was observed at 4 × magnification. Morphological changes were assessed in Col-0, *fer-4*, and *lacs2-3* leaf explants cultured on CIM for 0, 2, and 4 days. The distribution of de-methyl esterified pectin was visualized in Col-0 explants stained with Ruthenium Red at the same time points. In addition, Col-0 explants treated with DMTU were examined after 7 days of incubation on CIM to evaluate morphological effects.

### Image analysis

To measure the intensity and area of histochemical staining in images, Color Threshold function in ImageJ software (https://imagej.net/ij) was used.

### Illustrations and graphs

Bar graphs represent mean values, and whiskers indicate standard deviation of the mean (s.d.) or standard error of the mean (s.e.). For box plots, each box extends from the first quartile to the third quartile values, and center lines represent medium values. Whiskers indicate maximum and minimum values. Adobe Illustrator and Microsoft PowerPoint were used to generate illustrations.

### Statistical analysis

All statistical methods and the number of biological replicates in each assay are described in the figure legends. To determine statistically significant differences, one-way ANOVA followed by Tukey’s HSD test was performed using RStudio software, and Student’s *t*-test was conducted in Microsoft Excel. Bar graphs, box plots and heatmaps were generated using Excel. Asterisks and letter annotations in the figures indicate statistically significant differences at *P* < 0.05. Individual data points are shown in all plots in main figures.

## Notes

### Competing Interest Statement

The authors have declared no competing interest.

