## Supplementary Figures for "FERONIA defines intact tissue boundaries through cuticle development"

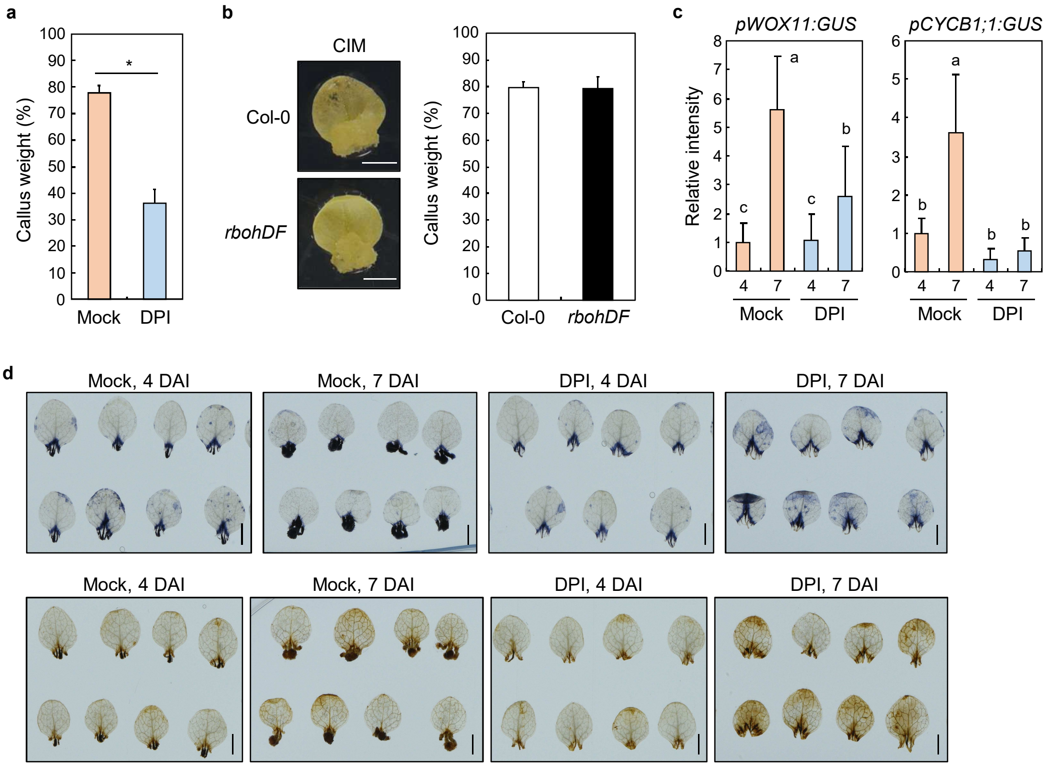


**Extended Data Fig. 1 | Effects of DPI on leaf callus formation.**

**a**, Callus weight in Col-0 leaf explants with or without DPI treatment. Callus weight (%) was calculated by dividing the weight of whole explants by that of the callus (*t*-test, $*P < 0.05$; $n = 3$). **b**, Mutation of *RBOHD* and *RBOHF* is not sufficient to alter callus formation. Leaf explants from Col-0 and *rbohDF* seedlings grown on MS-agar plates for 12 days were incubated on CIM. Callus formation was induced for 2 weeks in darkness. Percentage of callus weight was calculated by dividing the weight of whole plants by that of the callus. Two groups were statistically not significant (*t*-test, $P < 0.05$; $n = 4$). Scale bars, 0.25 cm. **c**, Quantitation of GUS intensity in Figure 1d. GUS intensity was measured using the ImageJ software. Different letters indicate statistically significant difference (Tukey's test, $P < 0.05$; $n$ = 15–16). X-axis indicates days after incubation (DAI). **d**, Additional replicates for Figure 1e. Several additional replicates for NBT (upper panel) and DAB (lower panel) staining for Figure 1e are displayed. Scale bars, 0.25 cm. For **a**–**c**, data represent mean ± s.d.

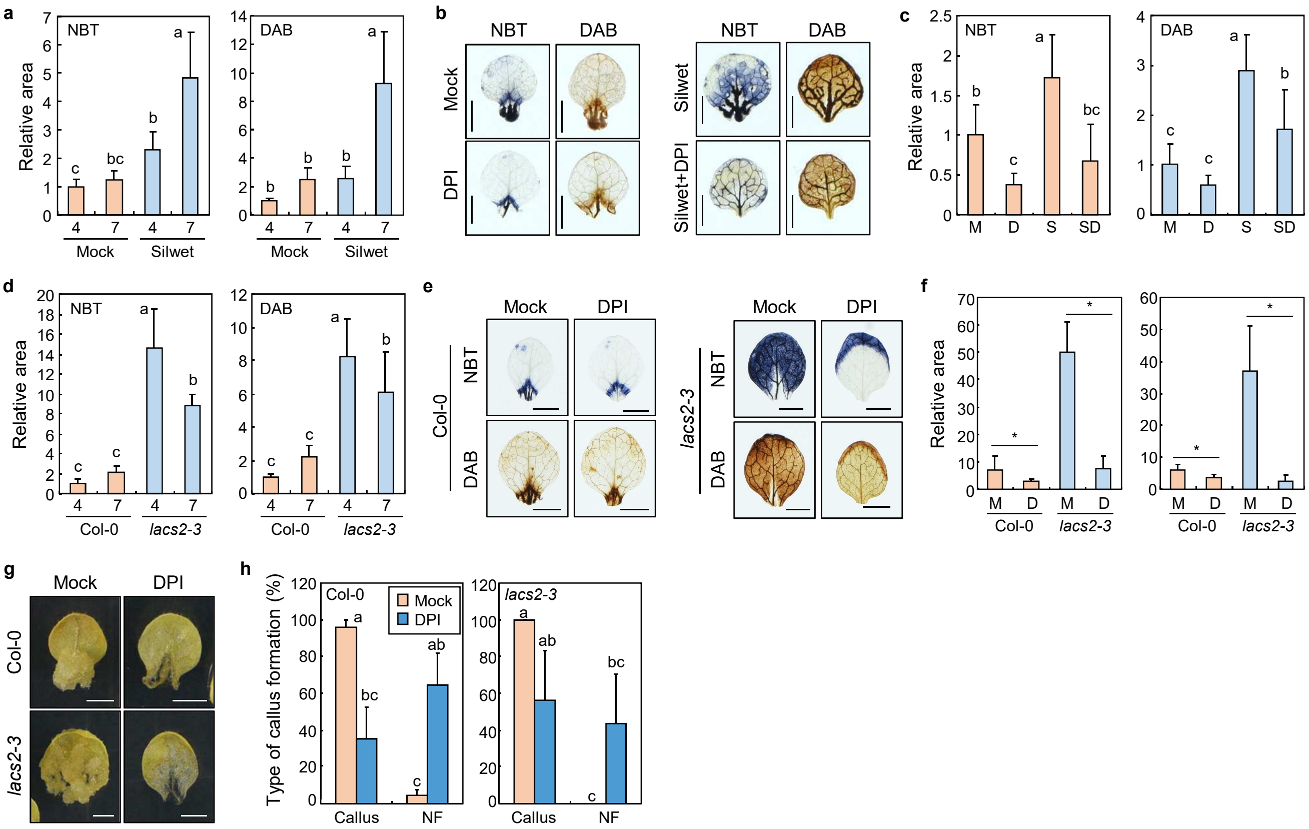


**Extended Data Fig. 2 | Relationship between cuticle permeability and callus formation.**
**a**, Quantitation of stained area for NBT and DAB in Figure 1h. Relative values are displayed. X-axis indicates DAI. Different letters indicate statistically significant difference (Tukey's test, $P < 0.05$; $n$ = 7–10). **b**,**c**, NBT and DAB staining of Col-0 leaf explants with or without DPI treatment. Representative staining images are displayed (**b**). Stained area for NBT and DAB was measured (**c**). M, Mock; D, DPI; S, Silwet; SD, Silwet+DPI. Different letters indicate statistically significant difference (Tukey's test, $P < 0.05$; $n$ = 7–13). **d**, Quantitation of stained area for NBT and DAB in Figure 1j. X-axis indicates DAI. Different letters indicate statistically significant difference (Tukey's test, $P < 0.05$; $n$ = 7–10). **e**,**f**, NBT and DAB staining of Col-0 and *lacs2-3* leaf explants with or without DPI treatment. Representative images are displayed (**e**). Stained area for NBT and DAB was measured (**f**). Statistical significance was determined by Student's *t*-test ($^{*}P < 0.05$; $n$ = 7–10). **g**,**h**, Callus formation in Col-0 and *lacs2-3* leaf explants with or without DPI treatment. Representative images of leaf callus are displayed (**g**). Percentage of callus-forming and non-forming leaf explants is displayed (**h**). Different letters indicate statistically significant difference (Tukey's test, $P < 0.05$; $n$ = 3). For **a**, **c**, **d**, and **f**, stained area was measured using the ImageJ software. For **a**, **c**, **d**, **f**, and **h**, data represent mean ± s.d. Scale bars, 0.25 cm (**b**, **e**, and **g**).

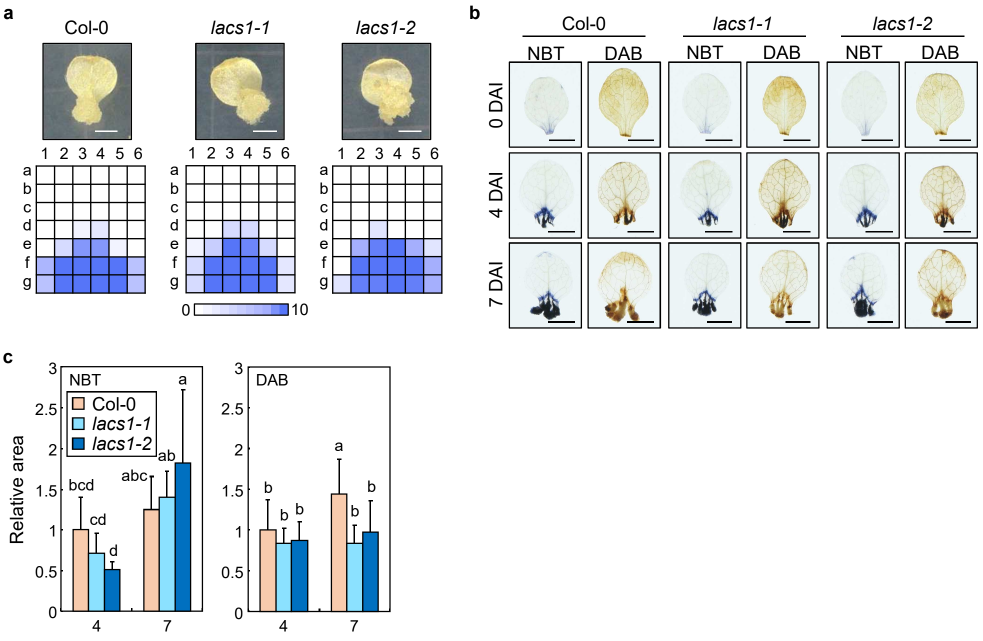


**Extended Data Fig. 3 | Callus formation in *lacs1* leaf explants.**
**a**, Callus formation *lacs1* mutants. Leaf explants from Col-0, *lacs1-1*, and *lacs1-2* seedlings grown on MS-agar plates for 12days were used for callus formation. Leaf explants were incubated on CIM for 2 weeks in darkness. Heatmap shows callus-forming sites. The number of explants forming callus at each site was visualized as a heatmap using 10 independent leaf explants. Scale bars, 0.25 cm. **b**,**c**, NBT and DAB staining in *lacs1* mutants. Leaf explants from Col-0, *lacs1-1*, and *lacs1-2* seedlings were incubated on CIM for the indicated time periods. Representative images are displayed (**b**). Scale bars, 0.25 cm. Stained area for NBT and DAB was measured using the ImageJ software (**c**). X-axis indicates DAI. Data represent mean ± s.d. Different letters indicate statistically significant difference (Tukey's test, $P < 0.05$; $n = 10$).

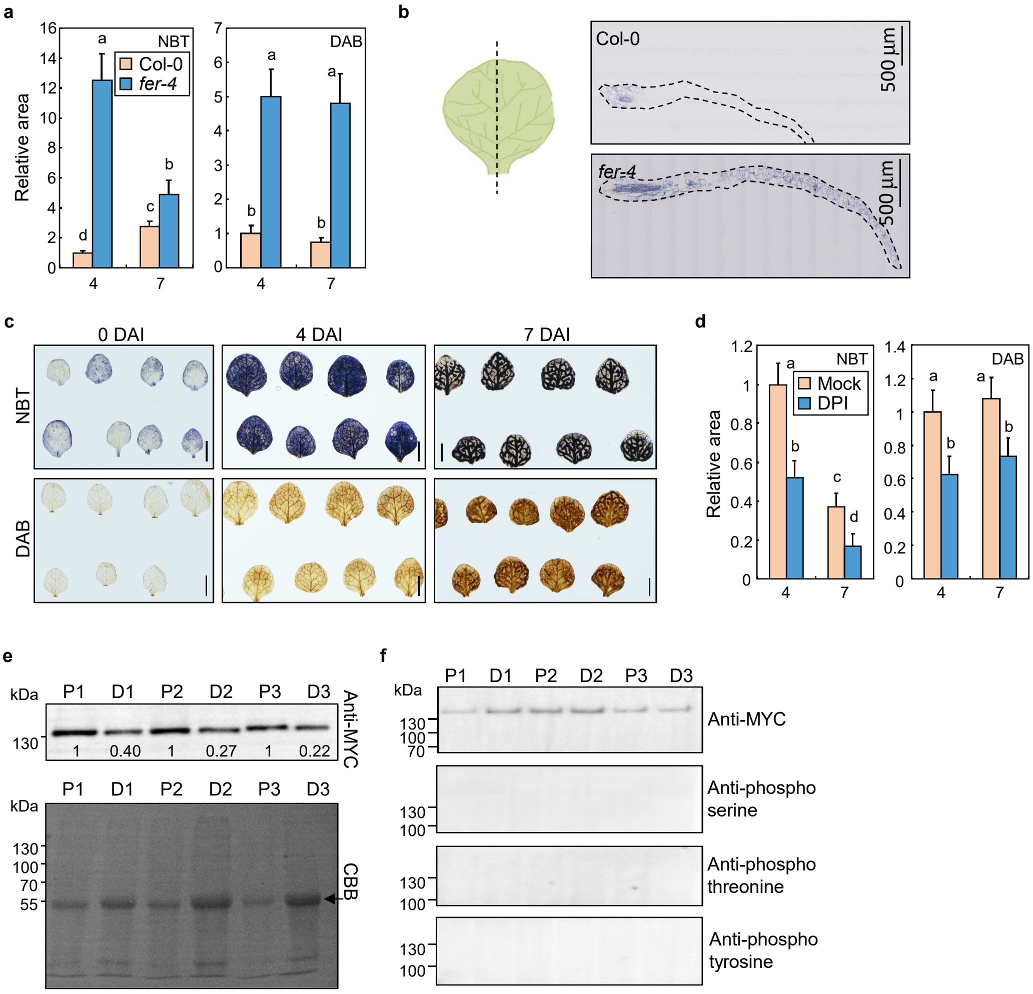


**Extended Data Fig. 4 | Regulation of spatial ROS distribution during callus formation in *fer-4*.** **a**, Quantitation of stained area for NBT and DAB in Figure 2b. Stained area was measured using the ImageJ software. X-axis indicates DAI. **b**, NBT staining of paraffin sections to detect superoxide anions in leaf explants at 2 DAI. Col-0 and *fer-4* leaf explants were stained with NBT and sectioned longitudinally (left panel). Dotted lines represent the outlines of sectioned tissues (right panel). **c**, Additional replicates for Figure 2c. Several additional replicates for NBT and DAB staining are displayed. Scale bars, 0.25 cm. **d**, Quantitation of stained area for NBT and DAB in Figure 2d. Stained area was measured using the ImageJ software. X-axis indicates DAI. **e**, Abundance of FER proteins in different parts of the explants during callus formation. *pFER:FER-MYC* leaf explants at 14 DAI were used. Proximal (P) and distal (D) regions of the explants were separately analyzed for FER-MYC protein abundance using immunoblot assays. Three biological replicates are displayed. Numbers indicate band intensities of FER-MYC proteins (anti-MYC), normalized to those of rubisco large subunit proteins [arrow, Coomassie brilliant blue (CBB)]. Band intensities were measured using the ImageJ software. **f**, Kinase activity of FER during callus formation. *pFER:FER-MYC* leaf explants at 14 DAI were used. FER-MYC proteins were collected by immunoprecipitation and then phosphorylated proteins were analyzed using phosphorylation-specific antibodies. Three biological replicates are displayed. For **a** and **c**, data represent mean ± s.d. Different letters indicate statistically significant difference (Tukey's test, $P < 0.05$; $n = 10$).

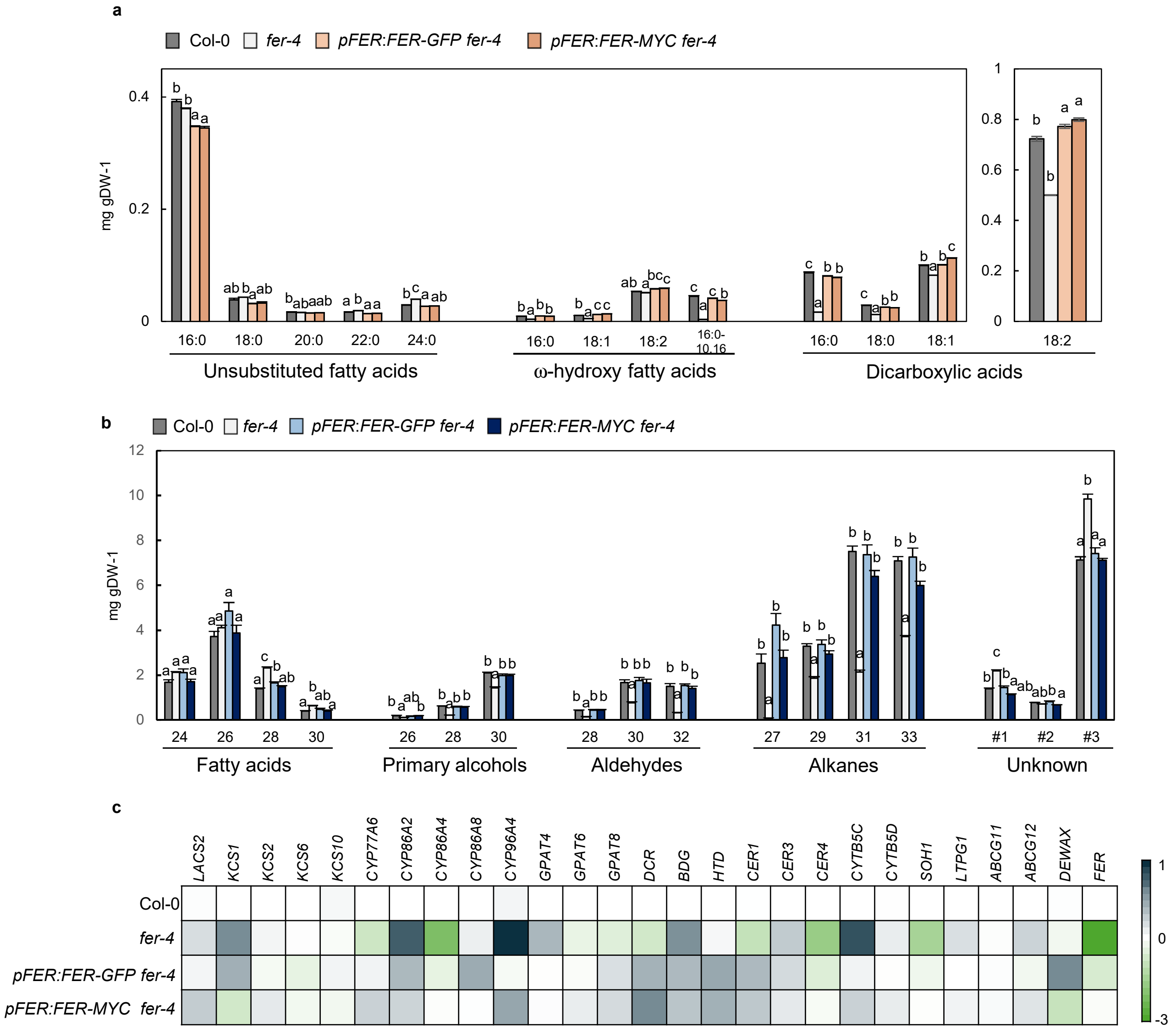


**Extended Data Fig. 5 | FER mediates cuticle formation in epidermal cells of aerial organs.** **a**,**b**, Total cutin (**a**) and wax (**b**) composition from the aerial organs of 12-day-old plants were analyzed using GC-FID. Data represent mean ± s.e. from five independent biological replicates. Different letters indicate statistically significant difference (Tukey's test, $P < 0.01$). **c**, Gene expression of cuticle biosynthesis-related genes in 12-day-old shoots was visualized as a heat map. Average expression values from three independent biological replicates are displayed as a heatmap Relative expression levels were converted to log2 scale. *PP2AA3* (At1g13320) was used to normalize gene expression.

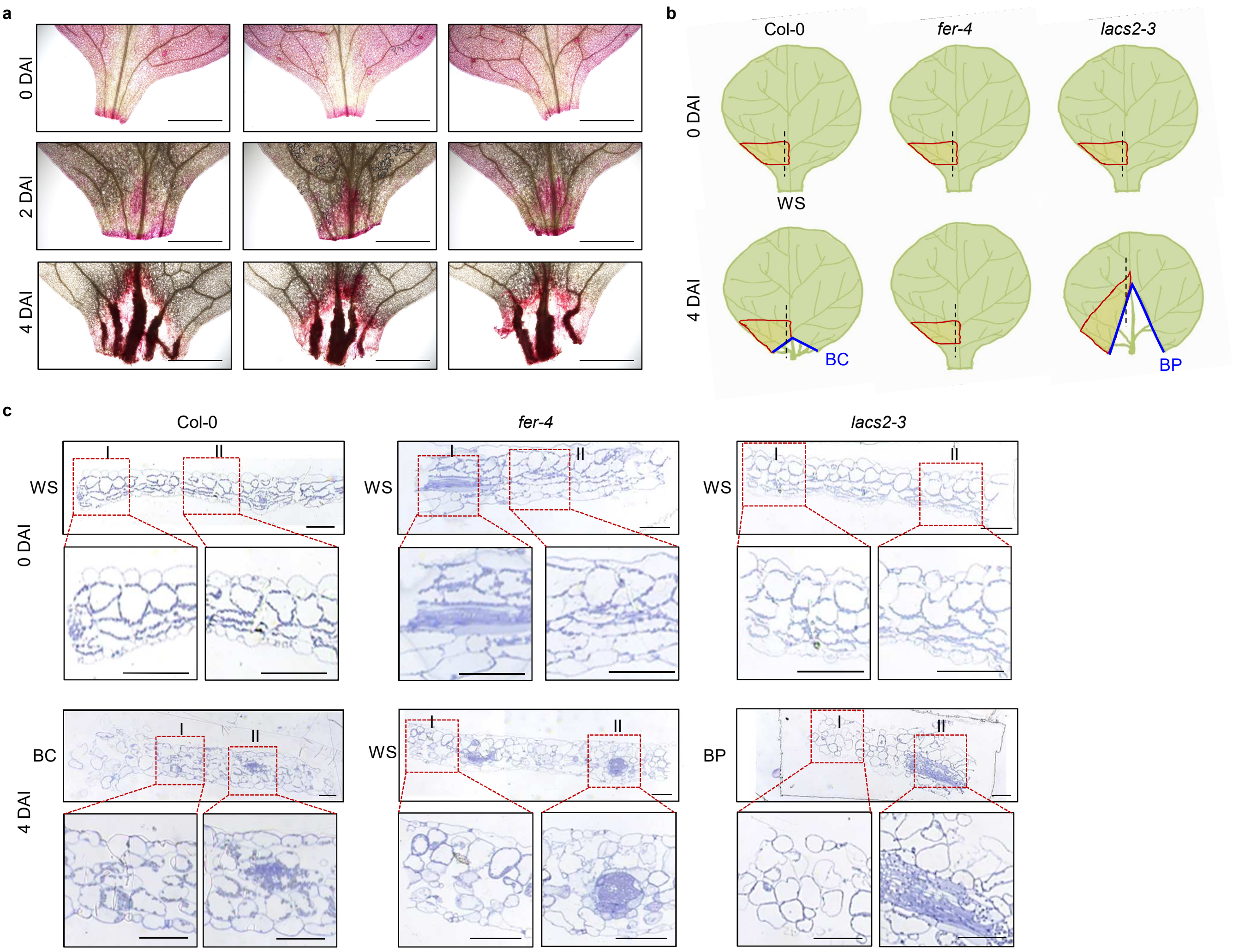


**Extended Data Fig. 6 | Imaging analysis near the BC of leaf explants.**
**a**, Additional replicates for Figure 4b. Several additional replicates for Ruthenium red staining for Figure 4b are displayed. Scale bars, 0.1cm. **b**,**c**, Sections of leaf explants during callus formation. Regions of leaf explants for TEM and light microscope imaging are illustrated (**b**). Longitudinal sections (dotted lines) within red-lined areas were subjected to light microscope (**c**) and TEM (Figure 4d) imaging after staining with 0.05% TBO solution and uranyl acetate/lead citrate solution, respectively. WS, wound site; BC, border of callus formation; BP, border of programmed cell death. Scale bars, 100 μm.

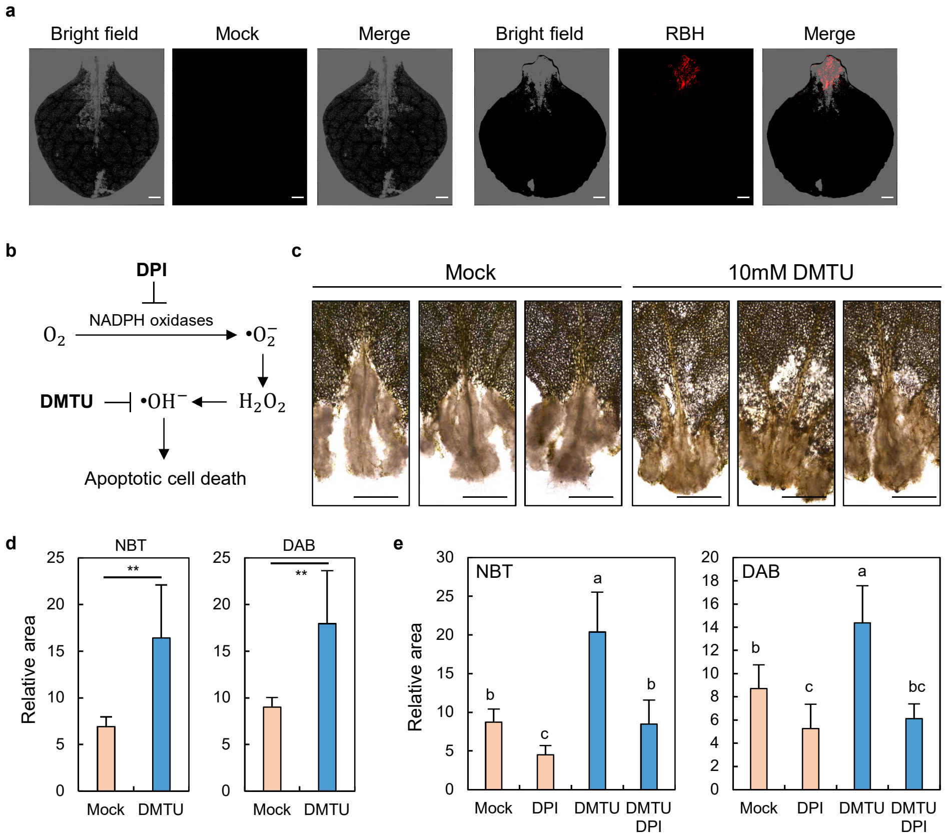


**Extended data Fig. 7 | Effects of DMTU on PCD and ROS distribution.**

**a,** Rhodamine B hydrazide (RBH) staining in Col-0. RBH staining was performed for hydroxyl radical imaging in leaf explants incubated on CIM at 3 DAI. Confocal microscopy was performed for bright field and RBH imaging. Mock indicates no RBH control. Scale bars, 0.05 cm. **b**, Pathways for the formation of superoxide anion, hydrogen peroxide, and hydroxyl radical. Inhibitors for specific ROS-forming pathways are shown. **c**, Additional replicates for optical microscopy in Figure 5a. Scale bars, 0.1 cm. **d**, Quantitation of stained area for NBT and DAB in Figure 5a. Stained area was measured using the ImageJ software. Data represent mean ± s.d. Statistical significance was determined by Student's *t*-test ($^{**}P < 0.01$; $n = 10$). **e**, Quantitation of stained area for NBT and DAB in Figure 5c. Stained area was measured using the ImageJ software. Data represent mean ± s.d. Different letters indicate statistically significant difference (Tukey's test, $P < 0.05$; $n = 8$–$10$).
